# Natural variation in the atypical resistance gene *Pita2* confers broad-spectrum neck blast resistance in rice

**DOI:** 10.64898/2026.01.31.702980

**Authors:** Ian Paul Navea, Mohammad Abul Monsur, Mary Jeanie Telebanco-Yanoria, Dianne Gene De La Rosa, Sherry Lou Hechanova, Arvin Paul Tuaño, Christian Joseph Cumagun, Il-Ryong Choi, Suresh Kadaru, Sung-Ryul Kim, Bo Zhou, Van Schepler-Luu

## Abstract

Neck blast (NB), caused by *Magnaporthe oryzae*, damages rice panicles and reduces yield. Knowledge of NB resistance remains limited due to the lack of reliable resistance evaluation methods. Here, we applied a newly established neck injection method and performed a GWAS on 335 diverse accessions from the 3K Rice Genomes Project to identify loci associated with NB resistance. We detected a significant association on chromosome 12, explaining 15-18% of the symptom variations caused by a highly virulent Philippine blast isolate (M64-1-3-9-1). Linkage disequilibrium analysis refined this region to a 42.3-kb interval containing *Pita2,* a known leaf blast resistance gene. We found that two *Pita2* allelic variants, *Pita2_a_* and *Pita2_c_*, both harboring the variant A/G (Lys879) in the last exon (Chr12:10,833,400), are associated with NB resistance. IR64 and a CO39 near-isogenic line (NIL) IRBLta2-Pi[CO] harboring *Pita2_a_* were resistant, whereas CRISPR-Cas9 knock-out of *Pita2_a_* in IR64 caused susceptibility to M64-1-3-9-1 and IK81-25. These results indicate that *Pita2_a_* is required for NB resistance. Furthermore, the CO39 NIL, IRBLta2-Pi[CO], and Lijiangxintuanheigu monogenic line (IRBLta2-Pi) harboring the *Pita2_a_* allele exhibited broad-spectrum resistance to 75% and 80% of Philippine differential blast isolates, respectively. The superior haplotype of *Pita2* contains two major SNPs (A/G and A/C at Chr12:10,833,400 and Chr12:10,845,095) occurs in 83% of IRRI elite breeding lines and can be used to select NB-resistant genotypes with an accuracy of 86%. Our findings identify *Pita2_a_* as a major gene for NB resistance and provide a valuable genetic resource for developing blast-resistant rice.

**PLAIN LANGUAGE SUMMARY:** Rice blast, caused by the fungus *Magnaporthe oryzae*, is a major threat to global rice production. Neck blast (NB) is the most severe type of blast, however, the genetic basis of NB resistance remains poorly understood. In this study, we analyzed 335 rice accessions to identify genes underlying the resistance against a Philippine blast isolate. We found that allelic variants *Pita2_a_* and *Pita2c* are strongly-associated with NB resistance. Knock-out of *Pita2_a_* allele made resistant rice plants susceptible while introgression into susceptible rice lines enhanced resistance to multiple blast isolates, confirming its role in NB resistance. Importantly, the superior alleles of *Pita2* are already present in 83% of elite breeding lines and can be used to select NB-resistant genotypes with an accuracy of 86%. Our findings clarify the genetic control of NB resistance and offer new tools for protecting rice yields in blast-endemic regions.

## 1 INTRODUCTION

Rice blast, caused by *Magnaporthe oryzae*, represents one of the most serious constraints to rice production (Zhang et al., 2016). The disease is responsible for up to 30% of global rice losses, enough to feed 60 million people (Nalley et al., 2016). There are several types of rice blast including leaf blast, node blast, neck blast and panicle blast depending on where the pathogen attacks. Leaf and neck blast are the most common forms of the disease, with the pathogen typically infecting leaves during the vegetative stage and the neck during the reproductive and ripening stages. Neck infection, regarded as the most damaging form, causes necrosis, interferes with nutrient assimilation and water movement, leads to panicle breakage, and results in either sterile panicles when infection occurs before the milk stage or poorly filled grains when it occurs at later stages (Ou, 1985).

Over 100 leaf blast resistance genes/QTLs have been identified, however, it is unclear whether these genes can also provide resistance against NB. Nineteen leaf blast resistance genes encode nucleotide-binding site leucine-rich repeat (NBS-LRR) proteins that recognize pathogen effectors in a gene-for-gene manner and trigger downstream immune responses, including induction of pathogenesis-related (PR) (Cesari et al., 2013; Okuyama et al., 2011; Sharma et al., 2012). Others such as *Pita2*, *pi21*, *Pi-d2*, and *Bsr-d1*, encode non-NBS-LRR proteins that can activate defense pathways including PR genes such *OsPR1a*, *OsPR1b*, and *OsWRKY45* (Chen et al., 2006; Zhao et al., 2018a; Zhou et al., 2022). Among the leaf blast resistance genes, *Pita2* (*Ptr*, LOC_Os12g18729) confers broad-spectrum resistance. The gene colocalizes with *Pita* on chromosome 12 and encodes an armadillo-repeat protein (ARM) required for *AVR-Pita* recognition (Greenwood et al., 2024; Xiao et al., 2024). The extent to which leaf blast resistance genes contribute to NB resistance has not been investigated.

NB progresses into panicle blast over time. Field screening has identified several QTLs associated with panicle blast, including *qPBR10-1* (Wu et al., 2021), *qPb6-1* (Fang et al., 2019), *qPbh-11-1* and *qPbh-7-1* (Fang et al., 2016), *Pi-jnw1* (Wang et al., 2016), *qPbm11* (Ishihara et al., 2014), *qPBR1* (Jinlong et al., 2024; Wang et al., 2024). To date, four panicle blast resistance genes including *Pb1*, *Pb2*, *Pb3*, and *Pb4* have been cloned and functionally characterized, however, the effectiveness of *Pb1* depends on additional QTLs, whereas the presence of natural resistance alleles and associated markers of *Pb2*, *Pb3*, and *Pb4* in elite varieties has not been examined. Despite functional validation, it is unclear whether these panicle blast resistance genes can provide resistance against NB. Previous studies indicate that panicle blast activates defense pathways distinct from leaf blast. This includes cell wall reinforcement and reactive oxygen species formation. Pathogen gene expression may also differ between the two tissues during infection (Du et al., 2024). Although some quantitative trait loci (QTLs) overlap, tissue- and stage-specific responses suggest largely independent pathways (Kalia & Rathour, 2019).

Coupling GWAS for resistance gene discovery with CRISPR-Cas9 mediated gene editing for gene validation can fast-track the identification of new disease resistance genes. GWAS exploit natural variation and historical recombination in diverse populations providing high mapping resolution while reducing reliance on bi-parental populations (Nakano & Kobayashi, 2020). The 3K-RGP is particularly powerful because of its dense marker coverage and diverse genetic base (Mansueto et al., 2016). Using this panel, novel *Pita2* and *Pia* alleles linked to leaf blast resistance were identified (Greenwood et al., 2024). In addition, 68 loci for blast resistance (Fan et al., 2024) and multiple loci for bacterial blight (Lu et al., 2021) were detected. These studies highlight the utility of the 3K-RGP in disease resistance trait discovery, as its size and balanced structure enhance the power of mixed linear (MLM) and multi-locus models. Candidate genes identified through GWAS can be rapidly validated using clustered regularly interspaced short palindromic repeats (CRISPR)-Cas9-mediated knockout or knock-in approaches, which allow precise functional testing of alleles in the same genetic background (Chen et al., 2022; Hussain et al., 2024; Thomson et al., 2022). However, identifying NB resistance genes or QTLs through GWAS has not been possible due to challenges in establishing reliable and reproducible evaluation protocols on a large set of rice accessions.

Several neck blast inoculation methods have been described, including brushing spore suspensions; placing infected leaf fragments from *Paspalum dilatatum* or *Brachiaria mutica* on the neck; wrapping the neck with cotton soaked in suspension; injecting spore suspensions into the neck or sheath; using plastic rings filled with conidial suspension; spraying; and applying slices of sporulating culture directly onto the neck (Bhaskar et al., 2017; Ghatak et al., 2013; Koga et al., 2004; Korinsak et al., 2023; Singh et al., 2021). Despite these various approaches, no standard inoculation method and disease measurement scale are available, and none has been applied to screen a large set of rice accessions for QTL mapping.

Recently, we optimized the neck injection method by adding a step in which inoculated plants were covered with lightweight polyethylene bags for 24 hours after applying the conidial suspension to favor pathogen proliferation (Yanoria et al., 2025). This enabled us to screen a large number of plants under partially controlled greenhouse conditions. We used this newly established artificial inoculation method and NB measurement severity scale to conduct GWAS on the 335 rice accessions from the 3K-RGP, leading to the identification of a major NB resistance locus located on chromosome 12 harboring *Pita2* with two natural variants, *Pita2_a_* and *Pita2_c_*, highly associated with the NB resistance. The role of *Pita2_a_* allele in NB resistance was characterized using gene introgression and CRISPR-Cas9 mediated gene knock-out. To support the deployment of *Pita2_a_*, we identified gene-based markers and surveyed elite breeding lines for the presence of resistance allele, linking trait discovery directly to breeding applications.

## 2 MATERIALS AND METHODS

### 2.1 Plant materials

A subset of 335 rice accessions (Table S1) from an original panel of 500 genetically diverse accessions described in a genome-wide association study (GWAS) on leaf blast (Greenwood et al., 2024) was used. The panel was derived from the 3000 Rice Genomes Project (3K-RGP) by excluding accessions carrying major blast resistance genes (*Pi2/9*, *Pik*, and *Pi3/5*) through gene-specific marker screening, and by further stratifying accessions according to country of origin and genetic subgroup. The 335 accessions were evaluated for neck blast (NB) resistance under controlled inoculation assays. Lijiangxintuanheigu (LTH) and CO39 were included as susceptible controls.

For functional validation of the candidate gene (*Pita2_a_*), multiple genetic resources were used including the BC_6_-derived CO39 near-isogenic line (NIL), IRBLta2-Pi[CO] and the BC_1_-derived LTH monogenic line (ML), IRBLta2-Pi, both harboring the *Pita2_a_* allele originating from Pi4. Additionally, the CO39 NIL, IRBLta-Ya[CO], harboring *Pita*, was included to differentiate the gene expression dynamics and resistance spectrum of *Pita* and *Pita2_a_*. Five T_2_ CRISPR-Cas9 knock-out lines (ko-1 to ko-5) generated in IR64, likewise harboring the *Pita2_a_* allele were evaluated. To estimate the frequency of the superior haplotype in breeding materials, 43 accessions were randomly selected from a panel of 101 elite IRRI breeding lines (Table S2) and evaluated for their NB reaction.

### 2.2 Plant growth conditions

Ten germinated seeds per accession were sown in plastic trays containing soil collected from the International Rice Research Institute (IRRI) field station in Los Banos, Laguna, Philippines. At 17 days after sowing (DAS), seedlings were transplanted into individual plastic pots (23 cm height, 16 cm diameter) in the BG-05 greenhouse at IRRI (14.1735°N, 121.2594°E). Each pot was filled with 3.5 kg of soil and fertilized at transplanting with 3.6 g ammonium sulfate [(NH₄)₂SO₄, 14-0-0], followed by additional applications three weeks later and before booting. Plants were grown under natural light at 25-32 °C with regular watering and pest monitoring. At 10-12 days after flowering, yellowing leaves were removed, and plants were transferred to a climatized room one day before artificial inoculation and maintained at 24-26 °C and 60-70% relative humidity under natural light.

### 2.3 Fungal isolates

We previously tested the leaf blast virulence of the 20 Philippine standard differential isolates (Telebanco-Yanoria et al., 2008) on 25 *Pi* monogenic lines in LTH background (Kobayashi, 2007). Among these, M64-1-3-9-1 was one of the most virulent, causing susceptibility in approximately 50% of the lines, including those harboring *Pia, Pii, Pik, Pik-p, Pik-h, Pib, Pi1, Pi3, Pi7(t), Pikm, Pita*, and *Pi11*. This isolate was therefore selected for evaluation of the GWAS panel. For functional validation in MLs, NILs, and knock-out mutants, and superior haplotype frequency estimation in selected IRRI elite breeding lines, isolates M64-1-3-9-1 and IK81-25 were selected based on the presence of *AVR-Pita* (Selisana et al., 2017), the cognate effector of *Pita2* (Xiao et al., 2024).

### 2.4 Neck and leaf blast resistance evaluation

NB resistance evaluation was performed using our newly established neck injection protocol (Yanoria et al., 2025). Five plants with a sufficient number and size of panicles were selected, and five panicles were inoculated per plant. Inoculation was carried out 10-12 days after flowering by injecting 100-200 µL of conidial suspension (1 × 10⁵ conidia/ml) approximately 5 mm below the panicle neck/collar with a 1-mL syringe. The plants were covered with a polyethylene bag for 24 hours immediately after inoculation. At ten days post-inoculation (dpi), disease response was evaluated based on lesion length (LL, mm; longest lesion from the injection point), disease incidence (DI, %; proportion of panicles infected relative to the total inoculated), and disease severity index (DSI, %) calculated from severity scores (0-9 scale; Table S3) using the formula:

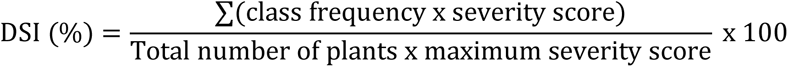

Lesion length was further used to classify accessions into four categories: resistant (R, 0-7 mm), moderately resistant (MR, >7-30 mm), moderately susceptible (MS, >30-55 mm), and susceptible (S, >55 mm). The GWAS panel and plant materials for functional validation were evaluated twice in a completely randomized design (CRD) with at least five biological replicates per accession.

Leaf blast resistance evaluation was performed using the IRRI standard protocol (Yanoria et al., 2025). Briefly, a tray of 14-day old seedlings was sprayed with 40 ml of the spore suspension (1 × 10⁵ conidia/ml) using an airbrush sprayer (0.5-mm nozzle atomizer, 18-20 psi, oil-free mini-compressor). Disease severity was evaluated at 6 dpi using the modified lesion type scale (0-2, resistant; 3-5, susceptible) (Yanoria et al., 2025). Leaf blast evaluation was performed twice.

### 2.5 Association mapping and candidate gene identification

For each rice accession in the GWAS panel, the data obtained from the two repeats were averaged to obtain the mean value for each phenotype. These accession-level means from the two repeats were used for downstream genome-wide association study (GWAS) analysis. Single nucleotide polymorphism (SNP) genotype data for the 335 rice accessions were obtained from the 3K-RGP dataset (https://snp-seek.irri.org/) and processed using PLINK v1.9 (Alexandrov et al., 2015; Purcell et al., 2007). From the 404k core SNPs in SNP-Seek, biallelic SNPs were extracted and filtered to remove minor allele frequency (MAF) of < 5%. Filtering resulted in a total of 253,465 SNPs for GWAS. All genomic coordinates were annotated against *Nipponbare* Reference Genome, IRGSP-1.0 (https://rice.uga.edu/). GWAS was conducted using an MLM (Zhang et al., 2021), in which SNP genotypes were treated as fixed effects, while population structure and kinship matrices were incorporated as random effects. Principal component analysis (PCA) for population structure estimation and kinship calculations was performed using TASSEL v5.0 (Bradbury et al., 2005).

Circular Manhattan plots were produced using the CMplot package in R (Yin et al., 2021). SNP density was visualized in 1 Mb bins. Genome-wide significance thresholds were calculated using a Bonferroni correction (P = 0.05/n, where n is the number of SNPs). A significance threshold was applied based on the Bonferroni correction (α = 0.05) on -log₁₀(P) values to identify marker-trait associations (MTAs). Quantitative trait loci (QTL) boundaries were defined by identifying SNPs in linkage disequilibrium (LD) with peak association signals and analyzing local LD patterns using PLINK v1.9 and Haploview v4.2, where candidate gene regions around peak SNPs were determined with the Solid Spine of LD function (Barrett et al., 2005).

### 2.6 Haplotype and candidate gene variant analysis

To identify the superior haplotype, SNPs located within the identified LD blocks were retrieved from the SNP-Seek database. Haplotype classification was guided by the SNPs spanning the LD blocks surpassing the Bonferroni significance threshold. The superior haplotype was defined as the one present in accession groups that exhibited resistance to NB across all three evaluated disease components. To assess the association between NB resistance and *Pita2* variants, all five known *Pita2* alleles (*Pita2_a_, Pita2_b_, Pita2_c_, Pita2_d_, and Pita2_e_*) were surveyed using the allelic sequence information published by Greenwood et al., 2024.

### 2.7 CRISPR-Cas9 construct design and rice transformation

A single guide RNA (gRNA) targeting the third exon of *Pita2_a_* (LOC_Os12g18729) was designed using CRISPR RGEN Tools (http://www.rgenome.net/about/). The gRNA sequence 5‘-GGCACACCTGCAGCCTGCGC-3’ (Protospacer adjacent motif, PAM: TGG) with the highest on-target score and minimal predicted off-targets was selected. Complementary single-stranded oligonucleotides encoding the 20 bp target sequence with *AarI* overhangs were synthesized as primers. The forward and reverse oligos were annealed and ligated into *AarI*-digested pSR339 (Figure S1), where gRNA expression was driven by the rice U3 small nuclear RNA (*OsU3*) promoter. The construct carries the *pZmUbi1-SpCas9-tNOS* cassette and a hygromycin resistance marker. The final construct was introduced into *Agrobacterium tumefaciens* strain LBA4404 using the freeze-thaw method (An et al., 1988). Rice transformation was performed in *Oryza sativa* ssp. indica cv. IR64 following an *Agrobacterium*-mediated transformation protocol established at IRRI (Slamet-Loedin et al., 2014). Of the 182 primary T0 transformants, 22 were selected for advancing to T1 based on the presence of predicted loss-of-function mutations. T_1_ plants were genotyped by target-region PCR amplification and Sanger sequencing. Five independent homozygous knock-out lines were advanced to T_2_ for phenotypic evaluation.

Additionally, we PCR-amplified and sequenced one potential off-target site that was predicted at Chr11:11,581,029, with three mismatches relative to the gRNA sequence (GGCACACCTGACAGCagGCGtAGG, with mismatches in lowercase). This site does not overlap with any annotated gene in the IR64 genome (https://riceome.hzau.edu.cn/). Primer sequences used for PCR amplification are listed in Table S4.

### 2.8 RNA extraction and qRT-PCR analysis

To quantify the expression of *Pita2_a_* and pathogenesis-related genes (*OsPR1b* and *OsPBZ1*), total RNA was extracted from neck tissues located approximately 20 mm below the last node of CO39, IRBLta2-Pi[CO] (*Pita2_a_*), and IRBLta-Ya[CO] (*Pita*) plants. Tissues were collected from M64-1-3-9-1 and mock-inoculated plants at 1-, 2-, 3-, and 5- dpi. Five biological replicates were used for each treatment and time point. RNA was extracted using TRIzol reagent (Thermo Fisher Scientific, Germany) according to the manufacturer’s instructions.

Complementary DNA was synthesized using the GoScript™ Reverse Transcriptase Kit (Promega, Madison, WI, USA). Quantitative real-time PCR was performed using Luna® Universal qPCR Master Mix (New England Biolabs, Ipswich, MA, USA) on a StepOne Real-Time PCR System (Applied Biosystems, Foster City, CA, USA) with gene-specific primers (Table S4). Gene expression levels were normalized to *OsActin1*, and relative expression was calculated using the 2^−ΔΔCt^ method (Livak & Schmittgen, 2001).

### 2.9 Statistical analysis

Differences among genotypes, allelic variant groups, and gene expression (*Pita2_a_*, *OsPR1b*, and *OsPBZ1*) were analyzed using one-way analysis of variance (ANOVA) implemented in the agricolae package in R (Jebesa & Kenea, 2023), followed by Duncan’s multiple range test (DMRT) for mean separation (Wanishsakpong et al., 2022). Reported P values were adjusted for multiple comparisons using either the Benjamini-Hochberg false discovery rate (Noble, 2009) or the Bonferroni method (Rastahi et al., 2024), with an adjusted P < 0.05 considered statistically significant. Pairwise comparisons were conducted using two-sample, two-tailed Welch’s *t*-tests (Curtis, 2024), applied both to contrast *Pita2_a_* and *Pita* monogenic lines across isolates and to evaluate resistance differences between LD-block haplotypes.

## 3 RESULTS

### 3.1 Rice accession from distinct subpopulations differ in resistance against neck blast

To assess neck blast (NB) responses and identify resistant donor lines, 335 diverse rice accessions from the 3K Rice Genome Project (3K-RGP) were inoculated with M64-1-3-9-1, a highly virulent *Magnaporthe oryzae* isolate from the Philippines. Three disease components including disease incidence (DI, %), disease severity index (DSI), and lesion length (LL, mm) were quantified. The panel comprised Indica (191), Tropical japonica (43), Aus (42), Subtropical japonica (21), Temperate japonica (18), Aromatic (11), and Admixed (11) accessions. In total, 26 accessions were resistant to M64-1-3-9-1 (Table S1, lines 1-26). We found that at the subpopulation level, Subtropical japonicas were the most resistant, showing the lowest DI, DSI, and LL values. Temperate japonica accessions were moderately resistant, while the Aromatic and Admixed groups showed intermediate resistance. The Indica group was generally susceptible, with longer LL values, whereas Aus and Tropical japonica were highly susceptible, displaying the highest DI and DSI (Figure 1; Table S5). These results demonstrate subpopulation-dependent differences in NB resistance.

**Figure 1.**
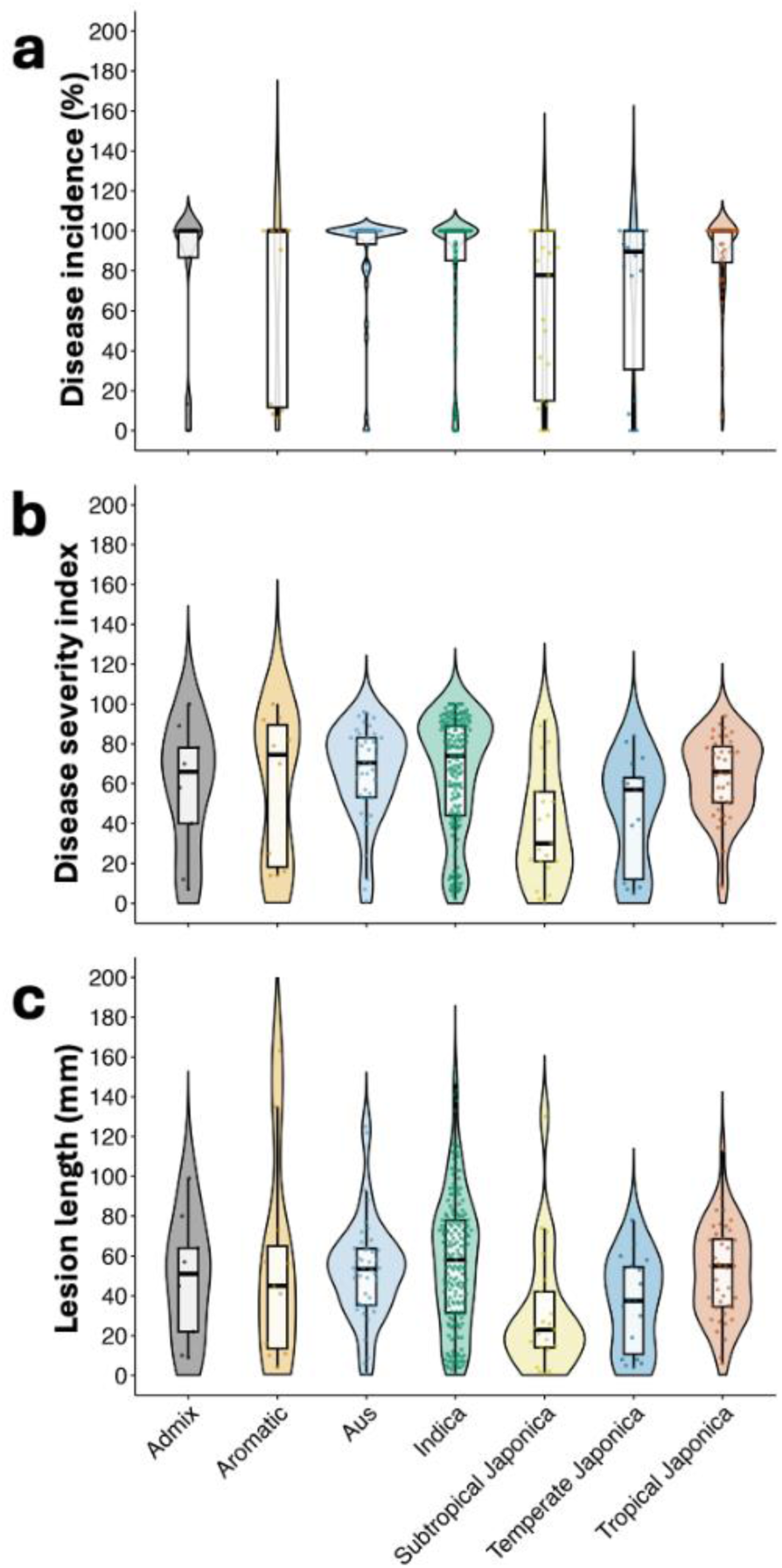
Subpopulation-level neck blast reaction in the 335 rice accessions. Violin plots show kernel-density estimates; overlaid thin boxplots mark the median (line), interquartile range (box), and 1.5×IQR whiskers; points are individual accessions. **(a)** disease incidence (%) **(b)** disease severity index, and **(c)** lesion length (mm).

### 3.2 Significant GWAS association for neck blast resistance maps to Pita2 locus

To identify loci associated with NB resistance, genome-wide association study (GWAS) was performed on 335 rice accessions for all three disease components: DI, DSI, and LL. A total of 253,465 single-nucleotide polymorphisms (SNPs), aligned to the *Nipponbare* reference genome, were retrieved from the SNP-Seek database. A strong association signal was consistently detected on chromosome 12 for all three traits at the Bonferroni-corrected threshold of −log₁₀(P) ≥ 6.7 (Figure 2a,b). We found that the SNP at Chr12:10,833,400 showed the strongest association with all three disease components, explaining 15%, 18%, and 14% of the phenotypic variation in DI, DSI, and LL, respectively. Fifty-six overlapping SNPs for DI, DSI, and LL between 10.58 and 10.92 Mb exceeded the significance threshold, defining a major-effect locus (Table S6). Linkage disequilibrium (LD) analysis using PLINK v1.9 and Haploview v4.2 refined this region to a 42.3-kb haploblock spanning 10.82-10.86 Mb and comprising three sub-blocks linked to the focal SNP (Figure 2c). This haploblock contains 19 additional SNPs besides the peak SNP (Table S7).

**Figure 2.**
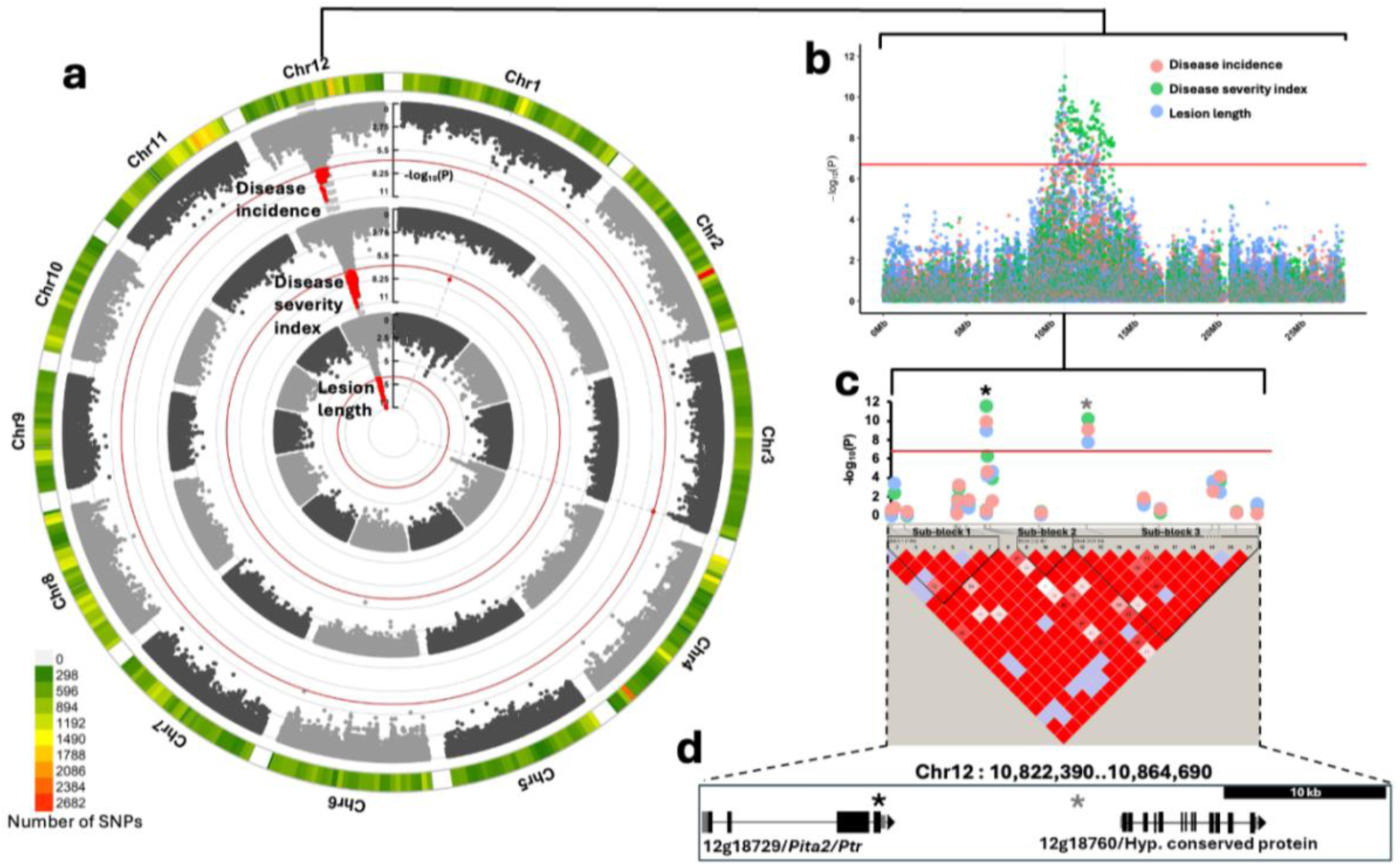
Genome-wide association analysis of neck blast resistance in rice. **(a)** Circular Manhattan plot depicting associations across all 12 chromosomes for disease incidence, disease severity index, and lesion length **(b)** Local Manhattan plot of chromosome 12 showing a major-effect locus centered on Chr12:10,833,400. **(c)** Linkage disequilibrium (LD) heatmap of the 42.3-kb interval (Chr12:10.82-10.86 Mb) highlighting the focal SNP and neighboring significant variants within the LD block. (**d**) Close-up view of the genomic region containing co-associated SNPs (haploblock) around the peak detected with isolate M64-1-3-9-1. The peak SNPs in the *Nipponbare* reference assembly are located at positions 10,833,400 (black asterisk) and 10,845,095 (gray asterisk). A 10-kb scale bar is shown in gray. Gene identifiers follow the Rice Genome Annotation Project (Osa1) Release 7 nomenclature, shortened by removing the LOC_Os prefix.

### 3.3 *Pita2_a_* and *Pita2_c_*, both carrying the Lys879 variant, are associated with NB resistance

To identify the candidate causal gene underlying the neck blast (NB) resistance association on chromosome 12, we examined the haploblock tightly linked to the peak association signal. This region harbored two annotated genes: LOC_Os12g18729 (*Pita2*, a known leaf blast resistance gene) and LOC_Os12g18760 (a hypothetical conserved protein) (Figure 2d).

*Pita2* encodes an armadillo-repeat protein (ARM) required for *AVR-Pita* recognition. Previous deep sequencing revealed five allelic variants (*Pita2_a_-Pita2_e_*) with polymorphisms confined to their C-termini, specifically the last 50 residues immediately downstream of the ARM domain (Greenwood et al., 2024; Xiao et al., 2024). Here, the peak SNP at Chr12:10,833,400 introduces a nonsynonymous G-to-A substitution in the last exon of *Pita2*, resulting in an arginine-to-lysine (Arg879Lys) amino acid change. To assess the functional relevance of this G/A variant, we examined its association with the NB resistance phenotype among the rice accessions. Of the 318 accessions carrying the G/A SNP, those with the A allele (Lys879) showed significantly lower DI, DSI, and LL than those with the G allele (Arg879) (Figure 3a). In addition, we surveyed this G/A SNP among the five *Pita2* alleles (*Pita2_a_*, *Pita2_b_*, *Pita2_c_*, *Pita2_d_*, and *Pita2_e_*). All accessions harboring the *Pita2_a_* or *Pita2_c_* allele had the Lys879 variant and exhibited significantly lower DI, DSI, and LL than those carrying Arg879 (Figure 3b-d; Table S8). These results indicate that *Pita2_a_* and *Pita2_c_*, both carrying the Lys879 variant, are associated with NB resistance.

**Figure 3.**
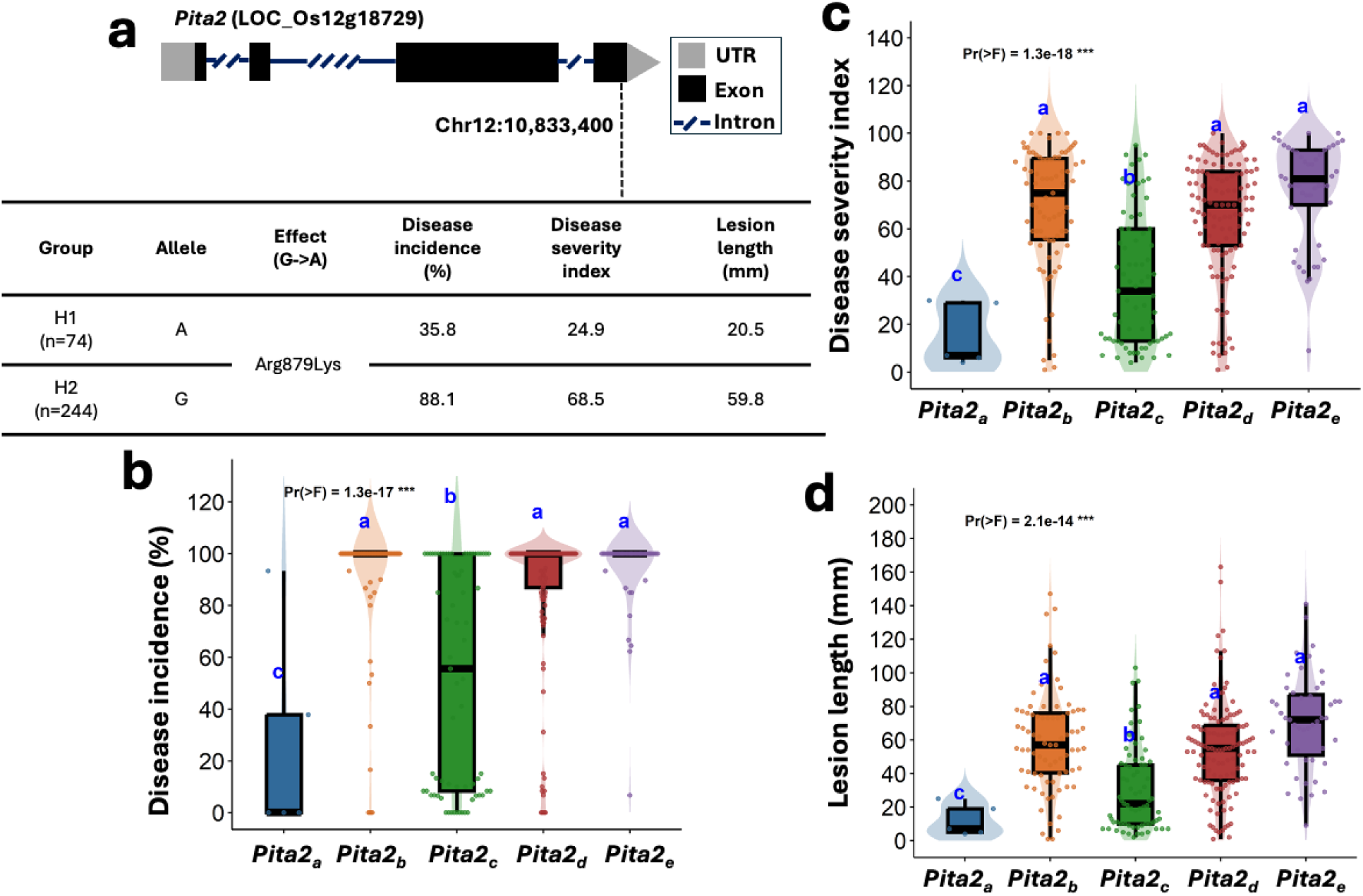
Neck blast responses associated with *Pita2* alleles. **(a)** Gene structure of *Pita2* (LOC_Os12g18729) showing the SNP at Chr12:10,833,400 in the last exon. The G to A substitution results in an Arg879Lys amino acid change. Phenotypic means for disease incidence, disease severity index, and lesion length are summarized for accessions carrying each allele. **(b-d)** Boxplots of disease incidence, disease severity index, and lesion length among five *Pita2* allelic variants. ANOVA results (Pr(>F) values) are shown above each panel, and different letters above boxplots indicate significant differences among allelic groups based on Duncan’s Multiple Range Test (DMRT, p<0.05).

### 3.4 *Pita2_a_* knock-out in IR64 led to susceptibility while its introgression in CO39 and LTH conferred broad-spectrum resistance to neck blast

To validate the function of *Pita2_a_* in NB resistance, we generated knock-out mutants in IR64, a NB-resistant variety harboring *Pita2_a_*, using CRISPR-Cas9-mediated gene editing. A single guide RNA was designed to target the third exon of *Pita2_a_* open reading frame (ORF). Five independent and homozygous T2 lines (*ko-1* to *ko-5*) with predicted loss-of-function mutations were selected and subjected to phenotypic analysis. Mutations in *ko-1*, *ko-2*, *ko-3*, and *ko-4* consisted of deletions of 49, 11, 5, and 6 bp, respectively, whereas *ko-5* carried single-nucleotide C insertion. All mutations resulted in frameshifts and premature stop codons, disrupting the *Pita2_a_* ORFs (Figure 4a,b). Phenotypic evaluation was performed by inoculating the knock-out lines and wild-type (WT) IR64 plants with two *M. oryzae* isolates, M64-1-3-9-1 and IK81-25, both known to be avirulent on *Pita2_a_*-containing lines (Table S9). All knock-out lines developed typical NB symptoms, with average lesion lengths ranging from 40 to 110 mm and 50 to 110 mm for M64-1-3-9-1 and IK81-25, respectively. IR64 WT plants showed no visible lesions under the same conditions (Figure 4c,d). To exclude off-target effects, we sequenced the predicted off-target site, which was intact in all edited lines (Figure S2), confirming that the observed susceptibility resulted from *Pita2_a_* disruption rather than off-target edits. These results demonstrate that *Pita2_a_* is required for IR64 NB resistance to both isolates.

**Figure 4.**
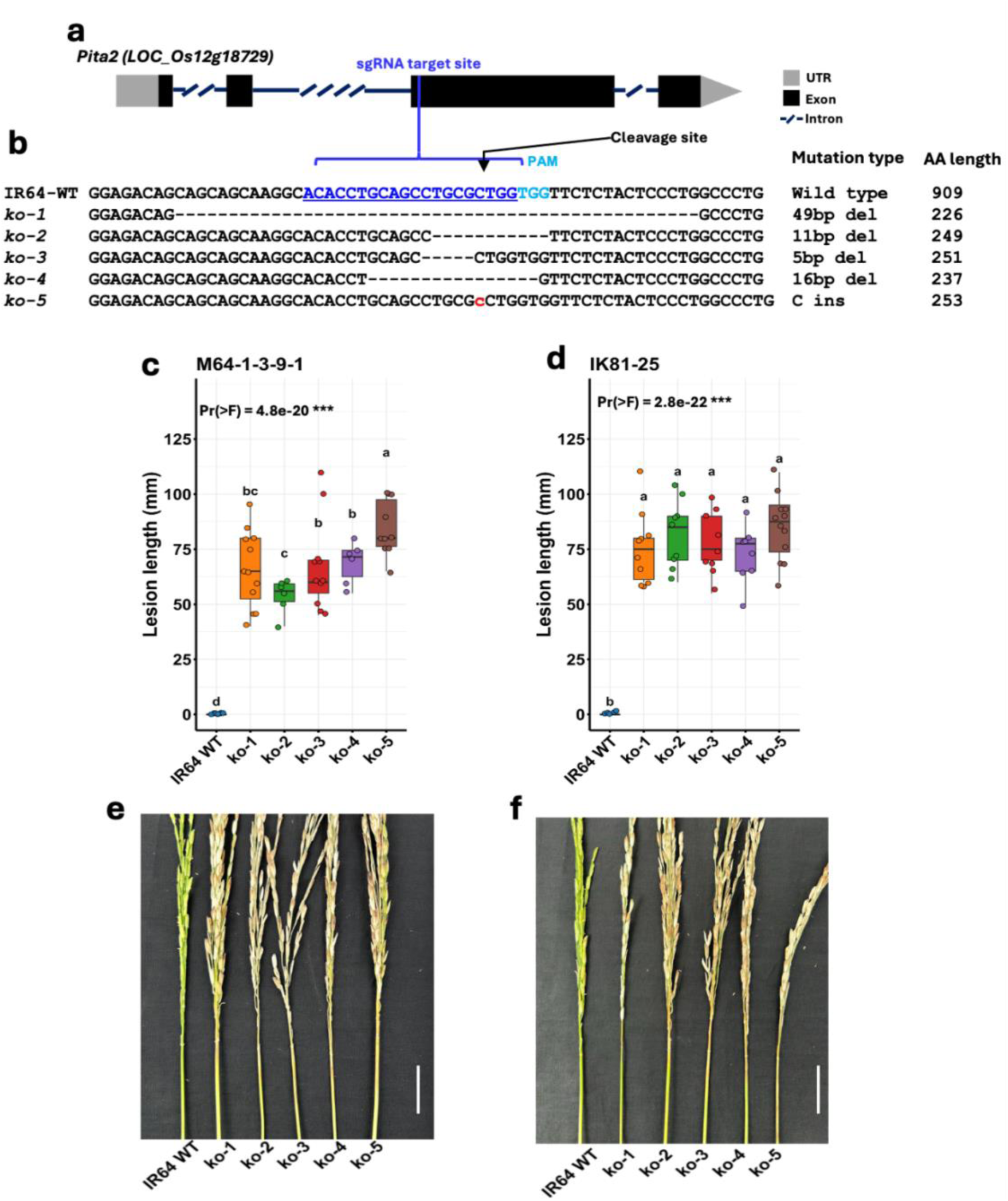
CRISPR-Cas9-mediated *Pita2_a_* knock-out in IR64. **(a)** Schematic representation of gRNA target sites within the genomic regions of *Pita2_a_*. **(b)** Mutation sites in knock-out mutants generated in the IR64 background via CRISPR-Cas9 editing **(c-d)** Boxplots lesion lengths in knock-out and WT plants following 10 days after inoculation with *M. oryzae* isolates M64-1-3-9-1 **(c)** and IK81-25 **(d)**. ANOVA results (Pr(>F) values) are shown above the boxplots, and different letters indicate significant differences among genotypes within an isolate based on Duncan’s Multiple Range Test (DMRT, p<0.05). **(e-f)** Representative panicles of the knock-out and IR64 WT plants inoculated with M64-1-3-9-1 **(e)** and IK81-25 **(f)**. White bars denote 20 mm.

To evaluate the NB resistance spectrum conferred by *Pita2_a_*, we used a Lijiangxintuanheigu (LTH) monogenic line (IRBLta2-Pi) and a CO39 near-isogenic line (IRBLta2-Pi[CO]) both harboring the Pi4-derived *Pita2_a_* allele (Kobayashi, 2007.). Twenty differential blast isolates representing the Philippine *M. oryzae* diversity were inoculated onto IRBLta2-Pi, while eight representative isolates were tested on IRBLta2-Pi[CO]. We found that IRBLta2-Pi was moderately resistant to resistant against 17 of the 20 isolates, whereas *Pita2_a_* introgression in CO39 (IRBLta2-Pi[CO]) conferred resistance to six of the eight isolates tested (Table S10). These findings indicate that *Pita2_a_* confers broad-spectrum resistance to NB. Furthermore, the IRBLta2-Pi displayed the same spectrum of resistance against leaf blast (Table S10), demonstrating the role of *Pita2_a_* in both leaf and NB resistance. Because *Pita* and *Pita2* are located at the same locus, we assessed an LTH monogenic line carrying *Pita* (IRBLta-CP1) to distinguish their individual contributions. IRBLta-CP1 was resistant to only four of the 20 isolates, suggesting that *Pita* is not the major NB resistance gene (Table S9, Figure S3).

### 3.5 *Pita2_a_* is induced upon pathogen infection and activates pathogenesis-related (PR) genes

To dissect the NB resistance mechanism, we assessed the expression dynamics of *Pita2_a_* upon pathogen infection. We performed qRT-PCR on M64-1-3-9-1 and mock inoculated CO39 and its near-isogenic lines harboring *Pita2_a_* (IRBLta2-Pi[CO]) and *Pita* (IRBLta-Ya[CO]). We found that expression of *Pita2_a_* was strongly induced in IRBLta2-Pi[CO], peaking at 2 dpi with an approximately three-fold increase relative to the susceptible CO39 and IRBLta-Ya[CO], before declining by 5 dpi (Figure 5a). In contrast, *Pita* introgression (IRBLta-Ya[CO]) did not significantly alter expression levels compared with the recurrent parent. These results indicate that infection by M64-1-3-9-1 induces *Pita2_a_*.

**Figure 5.**
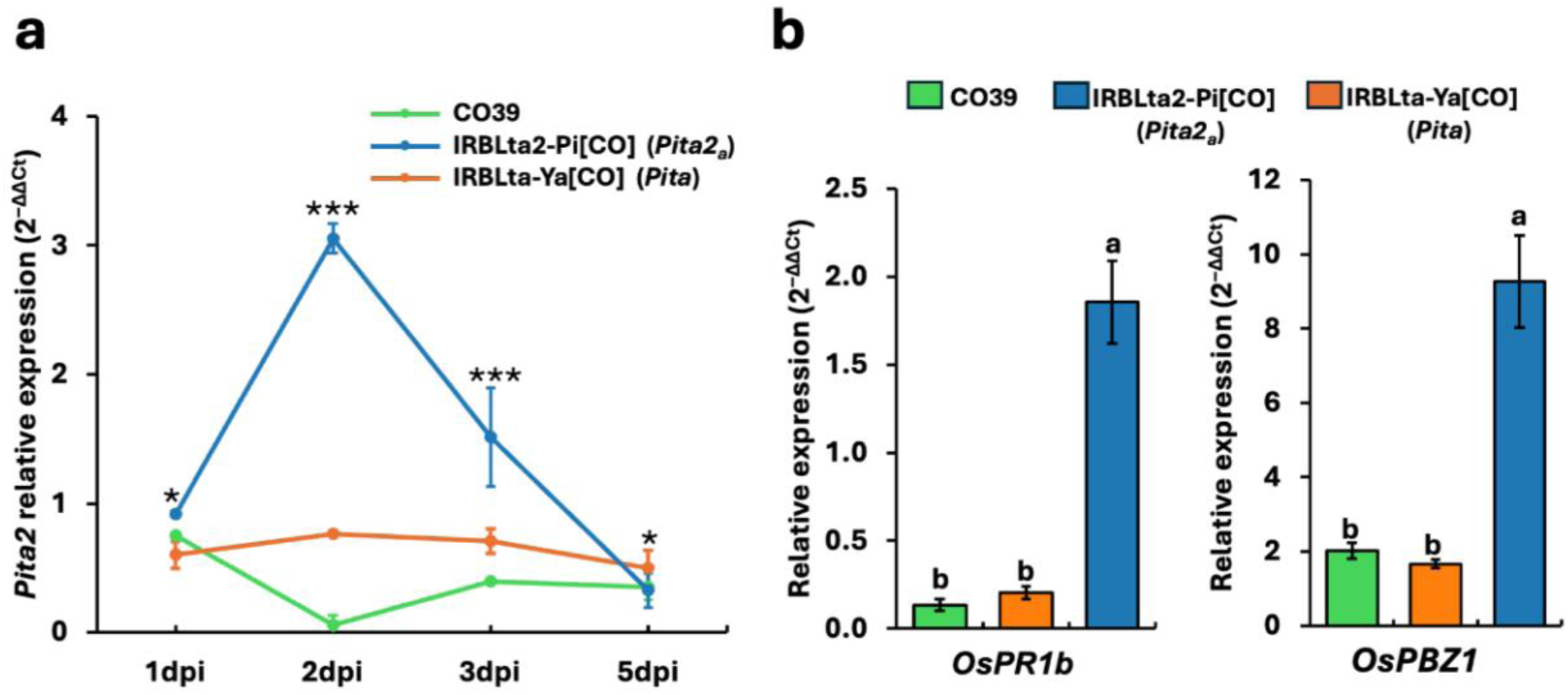
Expression dynamics of *Pita2_a_* and pathogenesis-related genes. **(a)** *Pita2_a_* transcript levels in CO39, IRBLta2-Pi[CO], and IRBLta-Ya[CO] at 1-, 2-, 3-, and 5-days post-inoculation (dpi). Expression was quantified by qRT-PCR using *OsActin1* as the internal reference. Asterisks indicate statistically significant differences among group means (*p<0.05, **p<0.01, ***p<0.001). **(b)** Relative expression of pathogenesis-related marker genes *OsPR1b* and *OsPBZ1* at 2 dpi. Bars represent mean ± SE of five biological replicates. Different letters above bars denote significant differences (P < 0.05; Duncan’s multiple range test).

To examine downstream defense activation of pathogenesis-related (PR) genes, we quantified transcript levels of two marker genes for effector-triggered immunity, *OsPR1b* (salicylic acid-mediated defense and systemic acquired resistance) and *OsPBZ1* (Probenazole-inducible gene 1; marker of broad-spectrum resistance). Both genes were markedly up-regulated in IRBLta2-Pi[CO] at 2 dpi but not in CO39 and IRBLta-Ya[CO] (Figure 5b). The enhanced expression of *Pita2_a_* and the downstream PR genes upon pathogen inoculation support the role of *Pita2_a_* as an active resistance gene that triggers defense signaling.

### 3.6 Superior haplotype of *Pita2* is rare in the 3K-RGP but enriched in IRRI breeding lines

To assess breeding relevance and develop SNP markers for selecting neck NB-resistant lines, we examined the distribution of the resistance-associated haplotype of *Pita2* among 2,132 accessions from the 3K-RGP Genebank and 168 IRRI elite breeding lines. Two peak SNPs within the LD block were used for haplotype classification: both exceeded the Bonferroni threshold for all disease traits and included the lead SNP at Chr12:10,833,400, which lies in the *Pita2* coding region, and a second significant SNP at Chr12:10,845,095 (Figure 2c, Table S7). Based on these SNPs, accessions were grouped into two haplotypes: H1 (AA/AA) and H2 (GG/CC) at Chr12:10,833,400/10,845,095. In the GWAS panel (335 accessions), 60 and 275 accessions belonged to H1 and H2, respectively (Figure 6a; Table 1). H1 was associated with resistance, showing significantly lower mean values for all disease metrics than H2 (Figure 6b); therefore, we designated H1 as the superior haplotype.

**Figure 6.**
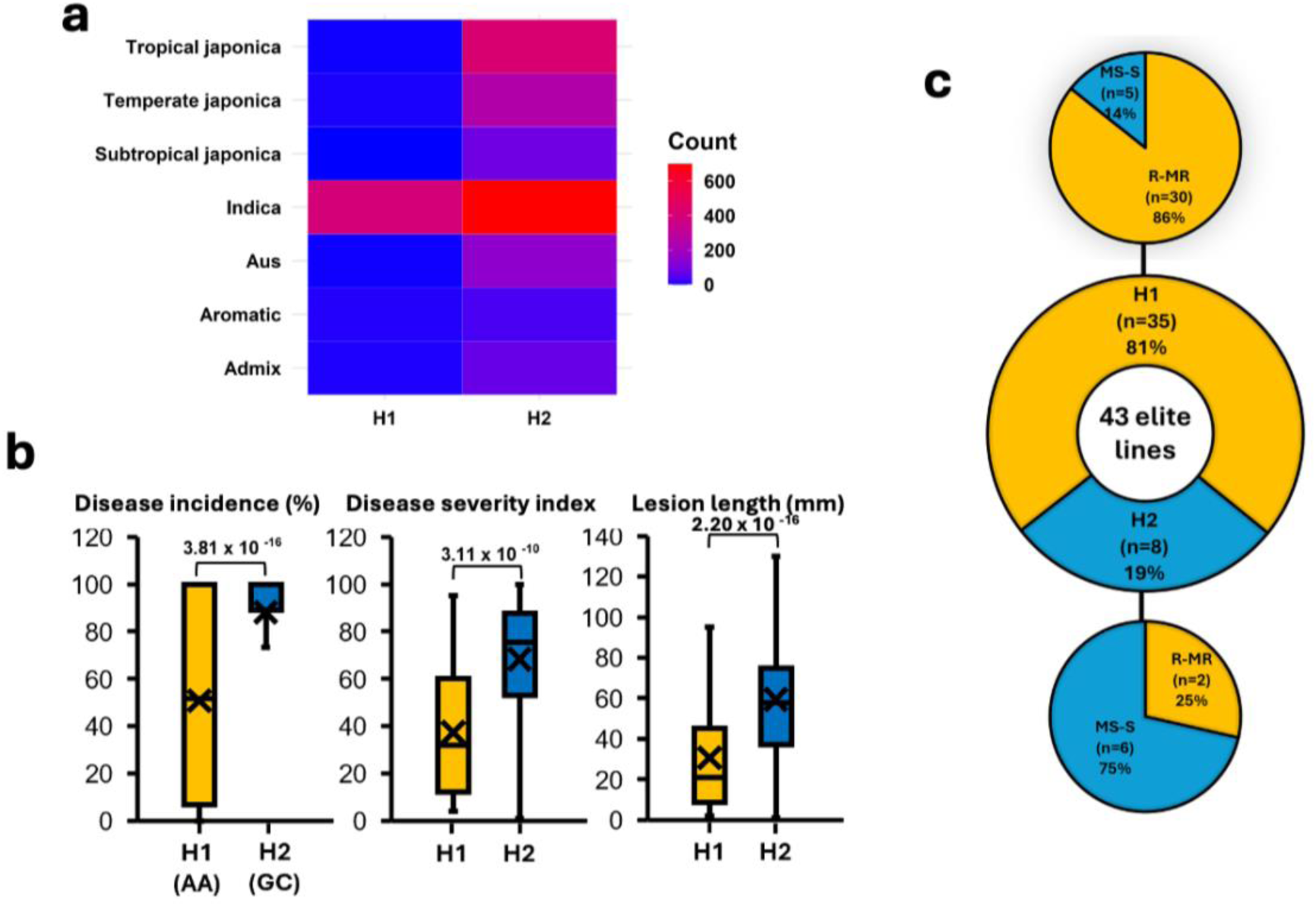
Distribution of haplotypes and their association with neck blast response. **(a)** Distribution of haplotypes (H1 and H2) across rice subpopulations. **(b)** Comparison of disease incidence, disease severity index, and lesion length between haplotypes. X marks the mean value **(c)** Neck blast reactions and haplotype abundance in the 42 IRRI elite breeding lines classified as resistant (R, ≤7 mm), moderately resistant (MR, >7-30 mm), moderately susceptible (MS, >30-55 mm), and susceptible (S, >55 mm) based on mean lesion length.

**Table 1.**
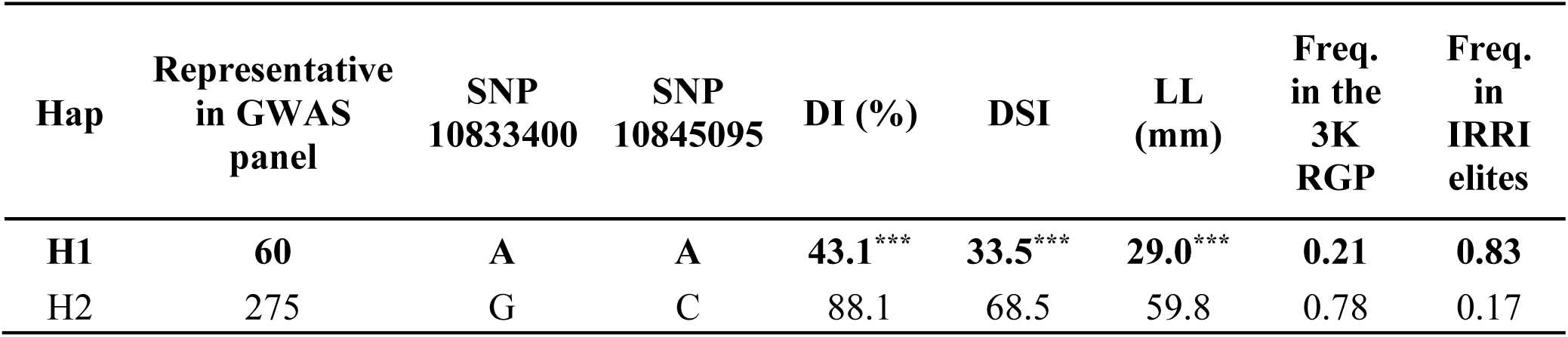
Haplotype analysis in the 3K rice genome panel (RGP) and IRRI elite breeding lines. Haplotype difference is summarized by asterisks (*** P<0.001) based two-sample t-test.

Filtering SNPs from the 3K-RGP classified 2,132 accessions into two groups, with 21% and 78% carrying H1 and H2, respectively, indicating that the superior haplotype is relatively rare among Genebank accessions. At the subpopulation level, the superior haplotype H1 is abundant in the Indica group (58%) but is absent from the Subtropical japonica group (Table S5). Among 168 IRRI elite breeding lines, 101 had complete genotype data for the two peak SNPs and were classified accordingly (Table S2). Of these, 83% carried the superior haplotype H1, indicating abundance of the resistant haplotype in IRRI elite germplasm.

To test whether H1 could serve as a selection marker for NB resistance, a random subset of 43 elite lines (Table S2, lines 1-43) was evaluated against the isolate M64-1-3-9-1. Among them, 35 lines carried H1, of which 30 lines (86%) showed resistance to moderately resistance (R-MR) reactions, while the remainder were moderately susceptible to susceptible (MS-S). In contrast, H2 carriers were predominantly MS-S (5/7; 71%), with only two lines (29%) classified as R-MR. Consequently, lines with H1 were approximately 15-fold more likely to be R-MR than those with H2 (odds ratio ≈ 15.0; based on 30/5 vs. 2/5) (Figure 6c). These results identify H1 as the superior haplotype strongly associated with NB resistance in the elite panel. Hence, H1 can be used to select NB-resistant genotypes with a positive predictive accuracy of 86% (Figure 6c).

## 4 DISCUSSION

In this study, we investigated the genetic basis of neck blast (NB) resistance in rice and identified *Pita2_a_* as a major gene underlying the resistance locus on chromosome 12. The Lys879 variant located in the C terminus of *Pita2_a_* was strongly associated with NB resistance. We also identified a superior haplotype (H1), defined by two SNPs (AA/AA at Chr12:10,833,400 and Chr12:10,845,095). H1 accurately predicts NB resistance with 86% and is prevalent among IRRI elite breeding lines. Consistent with earlier reports on the *Pita2_a_* locus’s broad-spectrum resistance, we found that this also extends to NB highlighting the utility of this allele in the improvement of multistage blast resistance in rice.

Through genome-wide association study (GWAS) of 335 rice accessions, we showed that the major locus for NB resistance maps to the *Pita2* region on chromosome 12. This association colocalized with signals previously reported for panicle blast in a pan-genome-wide association study using 414 accessions evaluated under natural field infection (McCouch et al., 2016; Wang et al., 2024). This indicated that natural variation in *Pita2* can modulate response to both panicle and neck blast. The concordance of these findings across datasets, environments, and population structures indicates the robustness and stability of the *Pita2* locus. A significant association for leaf blast resistance was also detected in this region (Greenwood et al., 2024), further demonstrating that *Pita2*-linked resistance spans multiple blast infection stages. Consistent with our results, the monogenic line harboring *Pita2_a_* (IRBLta2-Pi) exhibited the same resistance spectrum across 20 Philippines blast isolates in both leaf and neck blast (Table S10). Such cross-stage resistance is advantageous for breeding, as a single allele can enhance protection across leaf, panicle, and neck tissues.

The tight linkage between *Pita* and *Pita2* has historically complicated genetic dissection, because they are located in a low-recombination region near the chromosome 12 centromere (Wu et al., 2003). Although originally thought to be allelic (Rybka et al., 1997), subsequent analyses established that *Pita2* is distinct from *Pita* (Biswas et al., 2023; Meng et al., 2020; Zhao et al., 2018b). *Pita* encodes a nucleotide-binding site-leucine-rich repeat (NBS-LRR) protein (Bryan et al., 2000), whereas *Pita2* encodes an Armadillo repeat-containing (ARM) protein that confers broad-spectrum resistance against leaf blast (Zhao et al., 2018b). Although colocalized, the two genes act independently in mediating blast resistance (Meng et al., 2020; Xiao et al., 2024). In our results, *Pita2_a_* confers broader-spectrum resistance compared with *Pita* to both leaf and neck blast. The *Pita2_a_*-harboring monogenic line, IRBLta2-Pi was resistant to 16 of the 20 differential blast isolates from Philippines while the *Pita*-harboring IRBLta-CP1, was resistant to only four (Table S9, Figure S3). These results support the conclusion that *Pita2_a_* is a major contributor to NB resistance at the chromosome 12 locus.

Unlike the classical nucleotide-binding leucine-rich repeat (NLR) genes, which often show extensive allelic diversification due to strong selection pressures and frequent presence/absence polymorphisms (Yang et al., 2008), several non-NLR resistance genes in rice display comparatively limited natural variation. For example, *Xa4*, which encodes a wall-associated kinase (WAK), shows only a small number of haplotypes across surveyed germplasm panels (Edirisinghe et al., 2024). Similarly, the receptor-like kinase gene *Xa21* exhibits low allelic diversity in regional collections, with only a few haplotypes detected among dozens of cultivated varieties (Nanayakkara et al., 2018). *Pita2,* on the other hand is an example of non-NLR gene that possesses several alleles including *Pita2_a_*, *Pita2_b_*, *Pita2_c_*, *Pita2_d_*, and *Pita2_e_*, distinguished by amino acid substitutions and small Indels in their C terminus, specifically in the last 50 residues downstream of the ARM domain. Among these, *Pita2_a_*, which recognizes 11 known natural variants of *AVR-Pita*, lacks four residues at positions 870 to 873 (lysine-proline-glutamate-lysine) in its C terminus (Greenwood et al., 2024; Xiao et al. 2024). This four amino acid deletion is associated with broad-spectrum resistance to leaf blast. In this study, we identified a C terminus Lys879 variant consistently found among accessions harboring *Pita2_a_* and *Pita2_c_.* This Lys879 variant was strongly associated with NB resistance (Figure 3) suggesting that Lys879 can be a reliable marker for detecting *Pita2_a_* and *Pita2_c_* associated with NB resistance.

Although the C terminus of *Pita2* appears critical for resistance function, its mechanistic role in effector recognition remains unresolved. A recent study showed that *AVR-Pita* is the cognate effector of *Pita2* and that the *Pita2_a_* allele recognizes 11 natural *AVR-Pita* variants (Xiao et al., 2024). Since *Pita2_a_* and *Pita2_c_* carrying the Lys879 variant at the C terminus, are associated with NB resistance, we hypothesize that this residue influences the *AVR-Pita* binding affinity or recognition specificity. Further cloning and assays, e.g., yeast two-hybrid, co-IP, or cryo-EM are necessary to test this hypothesis. Understanding how *Pita2_a_* recognizes *AVR-Pita* will be critical for predicting resistance durability, given the rapid evolutionary turnover of *Magnaporthe oryzae* effector repertoires.

The *Pita2_a_* allele is expected to be effective across blast populations harboring *AVR-Pita* because it recognizes 11 known natural variants of this effector. Studies from Southern China, Vietnam, Philippines, and Southern India indicate that *AVR-Pita* occurs at relatively high frequencies at 72%, 58%, 57%, and 47%, respectively (Amoghavarsha et al., 2022; X. Chen et al., 2025; Lopez et al., 2019; Trang et al., 2019). This indicates the value of deploying *Pita2_a_* in these areas. Continued monitoring of *M. oryzae* populations is essential for targeted deployment. Coordinated evaluations across countries such as those led by the Global BioNET network established by IRRI would provide standardized data to guide durable deployment strategies and reduce the risk of resistance breakdown. Additionally, although we showed that *Pita2_a_* conferred broad-spectrum blast resistance in multiple genetic backgrounds, including CO39, Lijiangxintuanheigu (LTH), IR64, and selected IRRI elite breeding lines, validation across a wider set of varieties and agro-ecological zones is needed before large-scale deployment.

In the GWAS panel, only about 20% of the accessions carried the superior H1 haplotype, indicating that it remains relatively rare. Subpopulation differences in NB response were evident: Aus and Tropical japonica showed the highest susceptibility, Indica displayed intermediate responses, and Subtropical japonica the lowest disease scores. These subpopulation differences is consistent with the previous studies on leaf and panicle blast (Fan et al., 2024; Volante et al., 2020; Wang et al., 2024; Yu et al., 2022). Although H1 occurs frequently in Indica, the overall NB resistance in this group remains low likely because *Pita2* explains a modest 14 to 18% phenotypic variance. Additional loci within Indica likely contribute to susceptibility. Conversely, in Subtropical and Temperate japonica accessions, NB resistance was observed despite the low frequency of the superior haplotype, implying the presence of other resistance loci that may compensate for the absence of *Pita2_a_*. Population-specific analyses will be necessary to uncover these additional determinants of NB resistance.

The superior haplotype H1 is already enriched in elite irrigated breeding lines, providing an immediate resource for marker-assisted selection (MAS). In these lines, the high predictive accuracy (∼86%) of H1 likely reflects their more uniform genetic background, where locus effects are more consistent (Edwards et al., 2019; Isidro et al., 2015; Liu et al., 2023; Scutari et al., 2016). The high frequency of H1 in these lines suggests that breeders may have unintentionally selected for this haplotype during past improvement cycles. Such enrichment illustrates a broader pattern in rice breeding, where selection for agronomic traits can inadvertently alter the frequency of resistance alleles. For example, intensive selection for yield and quality traits reduced the prevalence of the *Piz*-*t/Pigm* allele in some breeding groups and were also observed for other blast-resistance loci (Gladieux et al., 2024; Thakur et al., 2013; Xiao et al., 2021). Similarly, analysis of NLR repertoires in rice detected signatures of balancing and positive selection on blast-related immune receptors such as RGA4 in Indica landraces, consistent with long-term selection maintaining resistance specificities (Gladieux et al., 2024).

However, H1 alone does not fully explain the phenotypic variation. Resistant H2 carrying lines demonstrate that the complementary loci also contribute to NB resistance. Thus, while H1 and key SNP markers (Table S7) are useful markers for selecting NB resistance, a multi-locus breeding strategy that integrates *Pita2_a_* with additional QTLs and incorporates genomic selection will be necessary for durable resistanc**e.** This is important because resistance conferred by single R genes can be rapidly overcome by pathogen evolution (Lore et al., 2025).

Although our validation confirmed that *Pita2_a_* is required for broad-spectrum resistance against the 20 Philippine isolates, the genetic architecture underlying NB resistance is possibly more complex. Several association signals surpassed the Bonferroni threshold (Table S6), indicating that additional loci may influence the phenotype. Genes that flank the major peak signal, in particular, may harbor variants that enhance or modify NB resistance together with *Pita2_a_*. For example, the peak association we detected in chromosome 12, aside from *Pita2*, is flanked by defense-related genes such as *Pita*, *Pi42(t)*, *OsPUB23*, *OsCYCA1*, *OsPR4*, and *OsPP130* (Figure S4). These genes could function in parallel defense pathways, contribute to quantitative resistance, or modulate the expression or effectiveness of *Pita2_a_*. Functional characterization of these candidates is therefore needed to determine whether they act independently, additively, or epistatically with *Pita2_a_* in shaping NB resistance.

The LD block we identified contained multiple SNPs in strong linkage with the peak association, any of which could serve as diagnostic markers for NB resistance. However, the predictive strength of individual SNPs may differ depending on populations structure and recombination history (Zhang et al., 2023). Mapping resolution is similarly influenced by historical recombination and LD patterns in the germplasm used (Padmashree et al., 2023). In rice, haplotype-based GWAS approaches can sometimes outperform single-SNP analyses, emphasizing that not all variants within an LD block have equal predictive value (Abed & Belzile, 2019). For breeding applications, prioritizing lead SNPs or a minimal set of highly informative markers will maximize efficiency while maintaining accuracy. The presence of multiple tightly linked signals also provides flexibility for marker design, allowing breeders to select SNPs compatible with their genotyping platforms without compromising reliability.

Defining the LD block therefore clarifies the genetic basis of resistance and provides a practical set of markers for breeding, allowing flexibility in the marker design across genotyping platforms in incorporating *Pita2*-based resistance into the breeding pipelines.

This study identifies *Pita2_a_* as the primary gene underlying broad-spectrum resistance to neck and leaf blast, with the Lys879 variant and superior H1 haplotype serving as predictors of resistance across diverse germplasm as well as in elite breeding lines. Our results clarify the genetic basis of the chromosome 12 resistance locus and provide markers suitable for accelerating the breeding of blast-resilient rice varieties. Beyond establishing the central role of *Pita2_a_*, our work illustrates the power of integrating genome-wide association, allele mining, and genome editing to uncover resistance mechanisms. The characterization of an effective non-NLR resistance gene highlights the contribution of noncanonical immune components and supports opportunities for precision breeding to enhance disease resilience. Collectively, these insights provide a strong foundation for the strategic deployment of durable blast resistance in global rice improvement programs.

## AUTHOR CONSTRIBUTIONS

VSL and BZ designed and initiated the project. IPN, MJY, VSL, BZ, IRC, CJC supervised the experiments. IPN, MAM, MJY, and APT performed the experiments. SRK and IRC designed the CRISPR-Cas9 construct. DGD and SLH made the construct and generated the CRISPR-Cas9 KO lines. SK selected and provided elite breeding materials. IPN and MAM analyzed the data.

IPN and VSL wrote the manuscript. All authors discussed the results, provided comments, and approved the final version.

## ACKNOWLEDGMENTS

We thank B. Fondevilla, C. Figuerra, and M. Apostol for their assistance in performing the phenotyping experiments. This work was supported, in whole or in part, by the Gates Foundation via the AGGRi project (INV-008226/ OPP1194925) and the CGIAR Research Initiative on Accelerated Breeding. The authors acknowledge the use of OpenAI’s ChatGPT (version 5.2) for minor language editing, including grammar and spelling.

## CONFLICT OF INTEREST

The authors declare no conflicts of interest.

---

**Table S1. Neck blast reactions of the 335 GWAS accessions inoculated with M64-1-3-9-1.** Values represent mean values of DI=disease incidence (%); DSI=disease severity index; LL=lesion length (mm).*Values represent lesion length (mm) and are classified as resistant (R, 0–7 mm), moderately resistant (MR, >7–30 mm), moderately susceptible (MS, >30–55 mm), or susceptible (S, >55 mm). nd=no data.

**Table S2. Neck blast reactions of IRRI elite breeding lines inoculated with M64-1-3-9-1.** Values represent lesion length (mm) and are classified as resistant (R, 0–7 mm), moderately resistant (MR, >7–30 mm), moderately susceptible (MS, >30–55 mm), or susceptible (S, >55 mm).

**Table S3. Modified Evaluation scale for neck blast disease (IRRI Plant Pathology Laboratory, 2019)**

**Table S4. Primers used in the qRT-PCR in the *Pita* and *Pita2* CO39 near-isogenic lines analysis and sgRNA off-target sequencing in CRISPR-Cas9 mutants.**

**Table S5. Subpopulation-level neck blast reactions of the 335 GWAS accessions inoculated with M64-1-3-9-1 and the distribution of peak SNPs haplotypes.** DI=disease incidence (%); DSI=disease severity index; LL=lesion length (mm). *values indicate number of accessions in the 3K RGP.

**Table S6. SNPs exceeding the Bonferroni-corrected genome-wide significance threshold.**

**Table S7. SNPs located within the 42.3-kb LD block on chromosome 12 (10.82–10.86 Mb) identified using Chr12:10,833,400 as the focal marker.** DI=disease incidence (%); DSI=disease severity index; LL=lesion length (mm).

**Table S8. Corresponding SNP 10,833,400 alleles and Pita2 alleles among the GWAS accessions used in this study.** Values represent mean values of DI=disease incidence (%); DSI=disease severity index; LL=lesion length (mm).

**Table S9. Neck blast reactions of the *Pita2* and *Pita* 25 LTH monogenic lines inoculated with the 20 Philippine standard blast isolates.** Values represent lesion length (mm) and are classified as resistant (R, 0–7 mm), moderately resistant (MR, >7–30 mm), moderately susceptible (MS, >30–55 mm), or susceptible (S, >55 mm). LTH was included as susceptible check.

**Table S10. Neck and leaf blast reactions of the monogenic line harboring *Pita2_a_*.** The line was inoculated with the 20 Philippine standard blast isolates. Values represent lesion length (mm) and are classified as resistant (R, 0–7 mm), moderately resistant (MR, >7–30 mm), moderately susceptible (MS, >30–55 mm), or susceptible (S, >55 mm). LTH was included as susceptible check for neck and leaf blast. ND = No data

**Figure S1. Map of the CRISPR-Cas9 binary vector pSR339 used in *Pita2_a_* editing.**

**Figure S2. *Pita2_a_* guide RNA off-target analysis. Off-target site sequences of the six *Pita2_a_* knock-out mutants aligned with the IR64 WT reference.**

**Figure S3. Neck blast reaction of IRBLta2-Pi (*Pita2_a_*) and IRBLta-CP1 (*Pita*) LTH monogenic lines (ML).** Lesion lengths of the monogenic lines across pathogen isolates. Bars show standard deviation of replicate observations, and overlaid points are individual replicates. Dashed horizontal lines at 7, 30, and 55 mm mark the lesion-length class boundaries used here: R (≤7 mm), MR (>7-30 mm), MS (>30-55 mm), and S (>55 mm). Within each isolate, significance of the *Pita2* vs *Pita* difference is summarized by asterisks (*p<0.05; ** p<0.01; **p<0.001; ns = not significant) based on a two-sample *t*-test.

**Figure S4. Defense-related genes flanking the peak association (Chr12:10,833,400) on chromosome 12.**

## Notes

### Competing Interest Statement

The authors have declared no competing interest.

