## Supplementary material for "Natural variation in the atypical resistance gene *Pita2* confers broad-spectrum neck blast resistance in rice": Supp. Fig

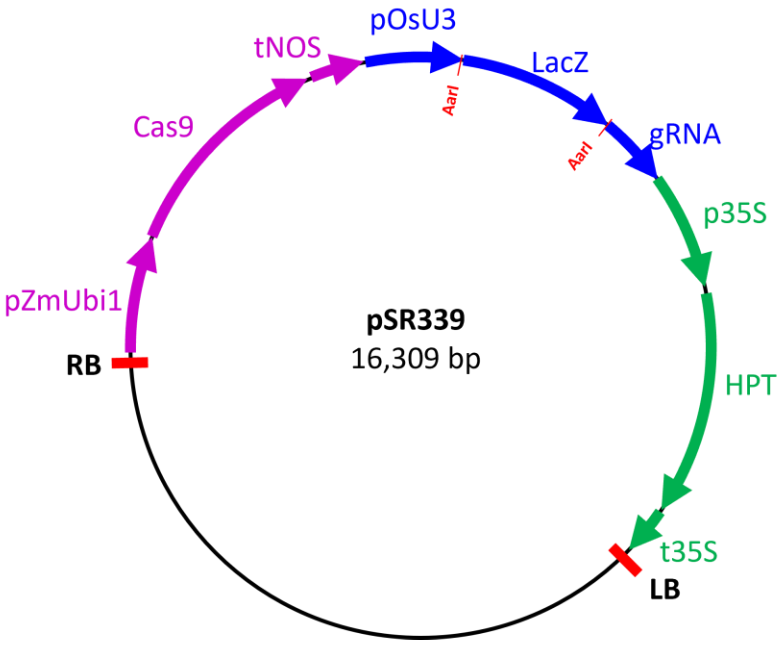


**Figure S1. Map of the CRISPR-Cas9 binary vector pSR339 used in *Pita2a* editing.**

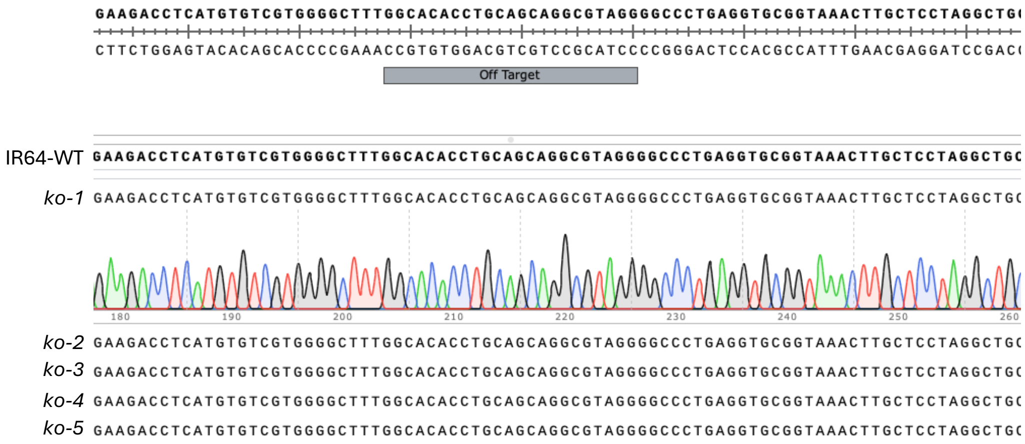


**Figure S2.** ***Pita2$_a$*** **guide RNA off-target analysis. Off-target site sequences of the six** ***Pita2$_a$*** **knock-out mutants aligned with the IR64 WT reference.**

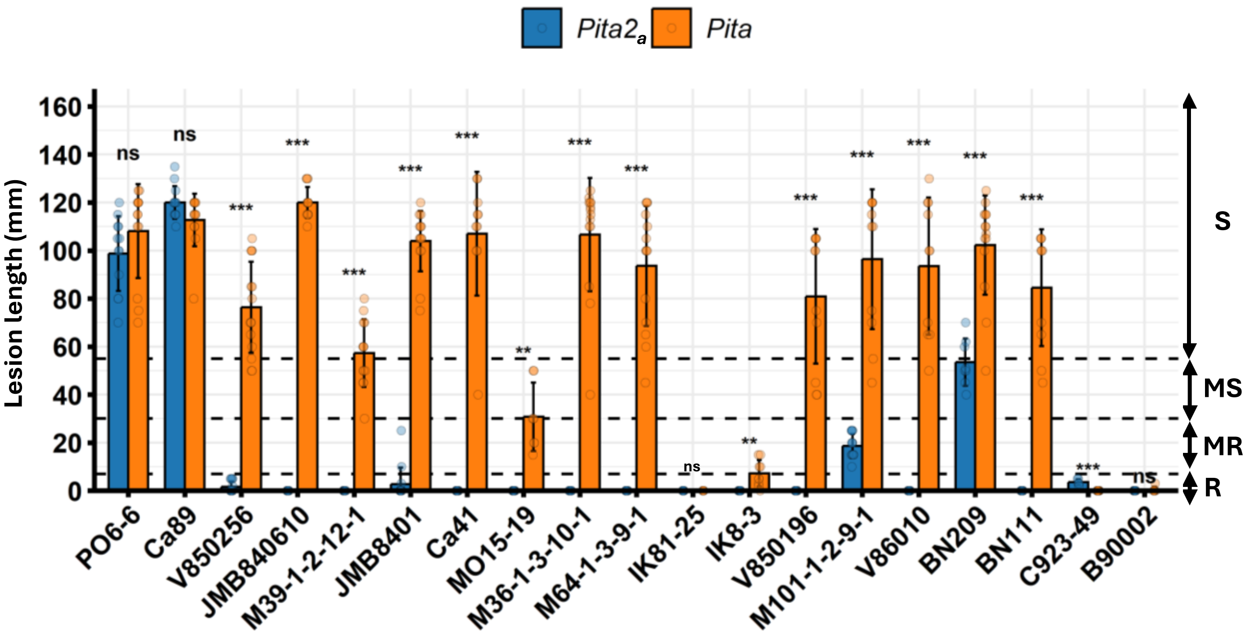


**Figure S3. Neck blast reaction of IRBLta2-Pi (*Pita2$_a$*) and IRBLta-CP1 (*Pita*) LTH monogenic lines (ML).** Lesion lengths of the monogenic lines across pathogen isolates. Bars show standard deviation of replicate observations, and overlaid points are individual replicates. Dashed horizontal lines at 7, 30, and 55 mm mark the lesion-length class boundaries used here: R (≤7 mm), MR (>7-30 mm), MS (>30-55 mm), and S (>55 mm). Within each isolate, significance of the *Pita2* vs *Pita* difference is summarized by asterisks (*p<0.05; ** p<0.01; **p<0.001; ns = not significant) based on a two-sample *t*-test.

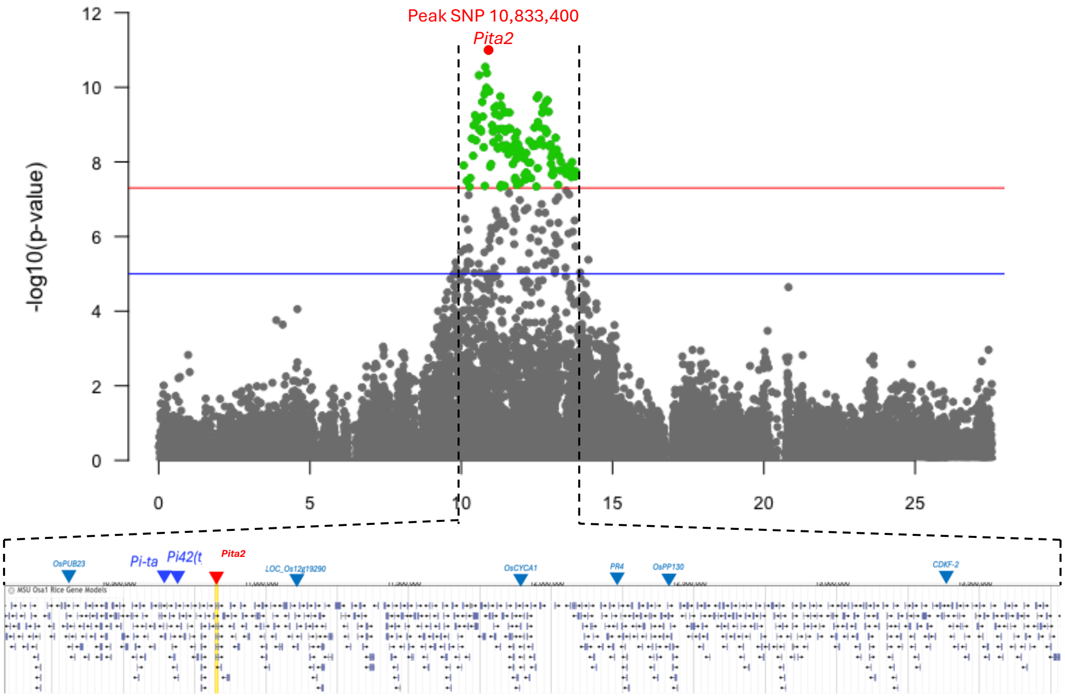


**Figure S4. Defense-related genes flanking the peak association (Chr12:10,833,400) on chromosome 12.**
